# DigiAra Computationally Designs Plant Mutants for Resistance to Microbial Infection in *Arabidopsis*

**DOI:** 10.64898/2026.08.07.743468

**Authors:** Tonghao Bai, Shuguang Cui, Yuning You

## Abstract

Plant breeding is a resource-intensive process that requires repeated cultivation and selection across multiple generations to develop varieties with desirable traits, yet computational tools capable of supporting this process remain limited. Here, we present DigiAra, an AI-based framework for designing *Arabidopsis thaliana* mutants with targeted traits, particularly enhanced microbial resistance. DigiAra implements an S3 pipeline—simulation, scoring, and screening: it simulates the transcriptional effects of genetic perturbations and microbial infections, scores the predicted responses in terms of relevant traits through biological pathway analysis, and screens candidate perturbations at multiple levels. In doing so, DigiAra enables the computational exploration of the genome-wide effects of genetic perturbations and diverse microbial infections in *Arabidopsis*. To develop DigiAra, we address two fundamental challenges. Methodologically, we introduce a hybrid architecture that integrates local gene-level interaction modeling with global transcriptional-state modeling to predict perturbation-induced changes in the *Arabidopsis* transcriptional state. From a data perspective, we establish a standardized pipeline for curating, harmonizing, and processing an integrated *Arabidopsis*–microbe transcriptional dataset comprising 495 samples from 26 projects. As a result, DigiAra accurately predicts gene-expression changes induced by unobserved genetic perturbations and microbial infections, achieving a Pearson correlation of 0.49. Moreover, it recapitulates the general non-self response (GNSR), a 24-gene program reflecting broad transcriptional reprogramming across bacterial perturbations. In an independent study, the predicted pattern-triggered immunity pathway scores further correlate with bacterial load, with a Pearson correlation of 0.57. Lastly, we deploy DigiAra to identify 27 gene knockouts through genome-wide screening that are predicted to enhance resistance to *Pseudomonas syringae* pv. tomato DC3000 (*Pst* DC3000) while limiting growth compromise, 9 of which are supported by published studies. Together, these results establish DigiAra as an effective framework for the computational design of *Arabidopsis* mutants. We have made our implementation openly available at https://github.com/youlab2025/DigiAra.

## Introduction

Plants constitute a substantial proportion of the biomass in Earth’s biosphere, and have played a crucial role in shaping human civilization by providing food, medicine, materials, and other essential resources [1]. The improvement of plants through cultivation and breeding has been a major human endeavor throughout history — plant breeding aims to develop varieties with desirable traits, such as resistance to specific diseases and tolerance to environmental stresses, including heat, drought, and nutrient limitation [2]. This is nevertheless challenging and requires considerable time and resources for several reasons. (i) Plant traits result from complex interactions among genetic and environmental factors [3], and these interactions are nonlinear, context-dependent, and difficult to model [4]. Moreover, plant breeding typically requires the simultaneous improvement of multiple traits, which may be interdependent or subject to physiological and genetic trade-offs [5, 6, 7]. (ii) Plant breeding involves a vast combinatorial search space [8]. Feasible genetic interventions and their combinations result in an enormous range of traits in different individuals. (iii) Evaluating each mutant or candidate line is costly. It requiring genetic manipulation, considerable cultivation resources, and repeated cycles of plant growth and evaluation [9, 10]. These challenges make the identification and development of suitable plant varieties a formidable search problem. To illustrate the scale of this challenge, developing and releasing a new crop variety through conventional breeding often spans several years and, in some breeding programmes, approximately a decade of repeated crossing, selection, and multi-environment evaluation [2, 11].

The resource-intensive nature of plant breeding motivates the development of computational models that can assist in the direct design and construction of plant mutants with desired traits, thereby potentially reducing the number of failed trials involving unsuitable mutants [12]. However, reliable models for this purpose remain lacking. Such models need to jointly capture the effects of intrinsic genetic variation and extrinsic disease or environmental conditions at the gene-expression level, which reflects system-wide plant states and provides informative readouts of plant traits [13]. The accumulation of RNA-sequencing datasets with condition annotations [14], together with advances in artificial intelligence (AI) techniques [15, 16], provides an unprecedented opportunity to develop models capable of predicting breeding outcomes. Nevertheless, the key challenge remains: how can external conditions be incorporated into representations of plant genetic states to model their joint effects on plant traits? Addressing this challenge could transform plant breeding from an empirical, trial-and-error process into a predictable, AI-guided design workflow.

Focusing on the model species *Arabidopsis thaliana* and the important trait of resistance to microbial infection, we present DigiAra, an AI-based framework for designing *Arabidopsis* mutants predicted to exhibit enhanced microbial resistance. DigiAra is built upon an S3 pipeline comprising simulation, scoring, and screening. First, DigiAra simulates the transcriptional responses of *Arabidopsis* to intrinsic, genome-wide genetic perturbations under extrinsic microbial infection conditions. Next, it scores different traits by analyzing their associated biological pathways based on the predicted transcriptional states. Finally, it screens candidate mutations for desired traits. From a biological perspective, DigiAra enables the computational exploration of the genome-wide effects of genetic mutations and diverse microbial infections on the transcriptional states of *Arabidopsis*.

To construct the simulation/prediction module of DigiAra, we address the methodological challenge of modeling interactions among the transcriptional state of *Arabidopsis*, genetic perturbations, and microbial infection conditions. Specifically, we develop a hybrid architecture that combines interaction modeling for local, gene-level interactions with interaction modeling between the global transcriptional state and experimental conditions. We further address the data challenge by developing a standardized pipeline to curate, harmonize, and process the first integrated *Arabidopsis*– microbe transcriptional dataset. The dataset currently comprises 495 samples collected from 26 projects and provides the foundation for developing DigiAra.

We demonstrate DigiAra’s predictive performance and its capability to design *Arabidopsis* mutants in experiments. For unobserved combinations of genetic perturbations and microbial infections, DigiAra accurately predicts transcriptional responses across all 27,448 genes, achieving a mean Pearson correlation of 0.49 with experimentally measured responses. In the screening task, DigiAra accurately ranks the predicted responses across conditions, with the resulting rankings exhibiting a mean Pearson correlation of 0.32 with those derived from experimental data. DigiAra’s predictive and screening capabilities are further supported by cross-dataset validation. By simulating the transcriptional response to the knockdown of each of the 27,448 genes under *Pseudomonas syringae* pv. tomato DC3000 (*Pst* DC3000) infection, DigiAra recapitulates a gene set that is significantly enriched in the predictions and has previously been identified as part of the general non-self response (GNSR), which is strongly associated with immune-related pathways. We further deploy DigiAra to design *Arabidopsis* mutants by jointly considering two objectives: immunity and growth. DigiAra reveals the global landscape of synergistic and antagonistic relationships between immunity and growth and identifies a set of candidate genes predicted to enhance immune responses while minimally suppressing growth. Among these candidates, 9 genes have previously been partially characterized for their roles in either immunity or growth.

## Results

### DigiAra implements a hybrid architecture to predict *Arabidopsis* genome-wide transcriptional responses to microbial infection and prioritize mutants for pathogen resistance

We develop DigiAra, an AI-based computational model that designs *Arabidopsis* mutants with enhanced resistance to microbial infection. DigiAra implements an S3 pipeline comprising simulation, scoring, and screening (Fig. 1C; Appendices A.4 and A.5). During simulation, it predicts the genome-wide transcriptional responses of *Arabidopsis* mutants to diverse microbial infections. During scoring, it evaluates the predicted differential expression of pathways associated with microbial resistance. During screening, it prioritizes mutants predicted to exhibit the strongest resistance. Through this pipeline, DigiAra systematically evaluates the effects of genetic perturbations in *Arabidopsis* and generates testable hypotheses for developing infection-resistant mutants.

**Figure 1:**
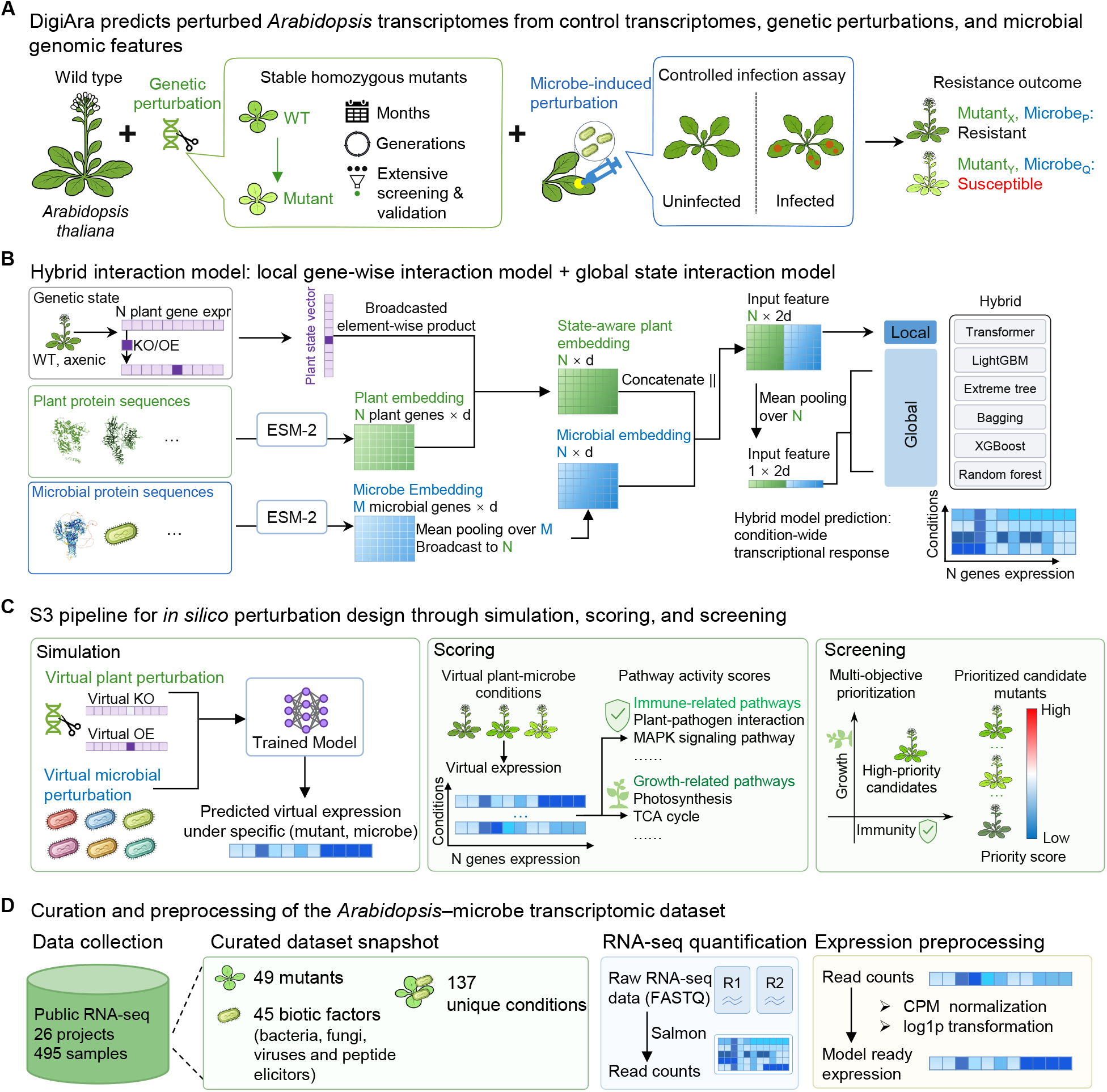
DigiAra implements a hybrid architecture to predict *Arabidopsis* genome-wide transcriptional responses to microbial infection and prioritize mutants for pathogen resistance. (A) Conventional identification of breeding-relevant plant variants requires the generation of stable homozygous mutants, controlled plant–microbe assays, and systematic assessment of plant responses across a vast combinatorial space of plant lines and microbial conditions, creating substantial bottlenecks in time and resources. (B) The DigiAra predictive module uses a hybrid interaction architecture to integrate the unperturbed plant transcriptional state, gene-wise genetic perturbation information, and ESM2-derived embeddings of plant and microbial proteins, thereby predicting condition-specific, genome-wide plant transcriptional responses. (C) The DigiAra simulation–scoring–screening (S3) pipeline designs and prioritizes candidate mutants predicted to exhibit desired immune–growth trait profiles. It first simulates plant-gene knockout or overexpression and microbial perturbations, alone or in combination, to predict condition-specific transcriptomes; then scores the predicted transcriptomes according to immune- and growth-related pathway activities; and finally screens and ranks candidate mutants through multi-objective prioritization. (D) Public RNA-seq data from 26 projects, comprising 495 samples, are curated into 49 plant genetic backgrounds, 45 biotic factors, and 137 unique conditions. Raw FASTQ reads are quantified with Salmon, followed by CPM normalization and log1p transformation to generate model-ready expression profiles.

Within the S3 pipeline, DigiAra’s simulation component relies on a predictive model that takes as inputs the transcriptional state of *Arabidopsis*, genetic perturbation information, and genomic features of the infecting microbe and predicts the resulting *Arabidopsis* transcriptional state under the combined perturbation and infection conditions (Fig. 1A). This machine-learning framework provides a computational analogue of the breeding process by modeling transitions in the plant’s molecular state across diverse genetic perturbations and infection conditions.

The DigiAra predictive module implements a hybrid interaction architecture that integrates the unperturbed transcriptional state, gene-wise mutant-state information, and ESM2-derived embeddings of *Arabidopsis* and microbial protein sequences to predict condition-specific, genome-wide transcriptional responses (Fig. 1B). Each condition is initialized using the corresponding unperturbed wild-type (WT) expression profile, represented by the WT–axenic state in Fig. 1B. To encode a gene knockout or overexpression, the expression value of the target gene is replaced with a predefined minimum or maximum value, respectively, while the expression values of all remaining genes are retained. This operation produces a gene-wise mutant-state vector that explicitly encodes the genetic perturbation while preserving the background transcriptional state. The *Arabidopsis* and microbial protein sequences are encoded separately using ESM2 [17]. Each *Arabidopsis* gene embedding is scaled by the corresponding value in the mutant-state vector, yielding a state-aware gene representation. The microbial protein embeddings are mean-pooled to obtain a microbe-level representation, which is then broadcast across all *Arabidopsis* genes and concatenated with each state-aware *Arabidopsis* gene embedding. Further details of the input encoding and feature construction are provided in Appendix A.2. The resulting gene-wise representations are provided to the gene-wise interaction model, whereas mean pooling across *Arabidopsis* genes generates condition-level features for the global interaction models. Predictions from the local and global models are subsequently hybridized through an ensemble strategy to generate a condition-specific, genome-wide *Arabidopsis* expression profile (Appendix A.4).

To provide the data foundation for DigiAra, we curate and process 495 publicly available RNA-seq samples from 26 independent projects (Fig. 1D). The resulting dataset comprises 49 plant genetic backgrounds, 45 biotic factors, and 137 unique experimental conditions. The biotic factors span bacteria, fungi, viruses, and peptide elicitors, enabling the predictive module to learn plant responses to diverse biotic challenges. Raw FASTQ reads are quantified using Salmon [18], followed by CPM normalization and log1p transformation to generate standardized, model-ready expression profiles (Appendix A.1). This integrated dataset provides the foundation for training the DigiAra predictive module, while the S3 pipeline translates its transcriptional predictions into trait-oriented virtual screening across a broad combinatorial space of *Arabidopsis* lines and microbial challenge conditions.

### DigiAra accurately predicts *Arabidopsis* transcriptional responses to unobserved genetic perturbations and microbial infection conditions

We ask how accurately DigiAra predicts *Arabidopsis* transcriptional responses under unseen genetic perturbation and microbial infection conditions. To assess this, mutant-only, microbe-only, and combined mutant–microbe conditions were partitioned at the condition level into training, validation, and test sets at a 7:2:1 ratio (Fig. 2A; Appendix A.3). DigiAra was trained on the training set, optimized using the validation set, and evaluated on the held-out test set. Performance was assessed using multiple metrics, including Pearson correlation, from two complementary perspectives: sample-wise evaluation, which compares predicted and observed genome-wide expression profiles within each sample, and gene-wise evaluation, which compares predicted and observed expression levels for each gene across samples.

**Figure 2:**
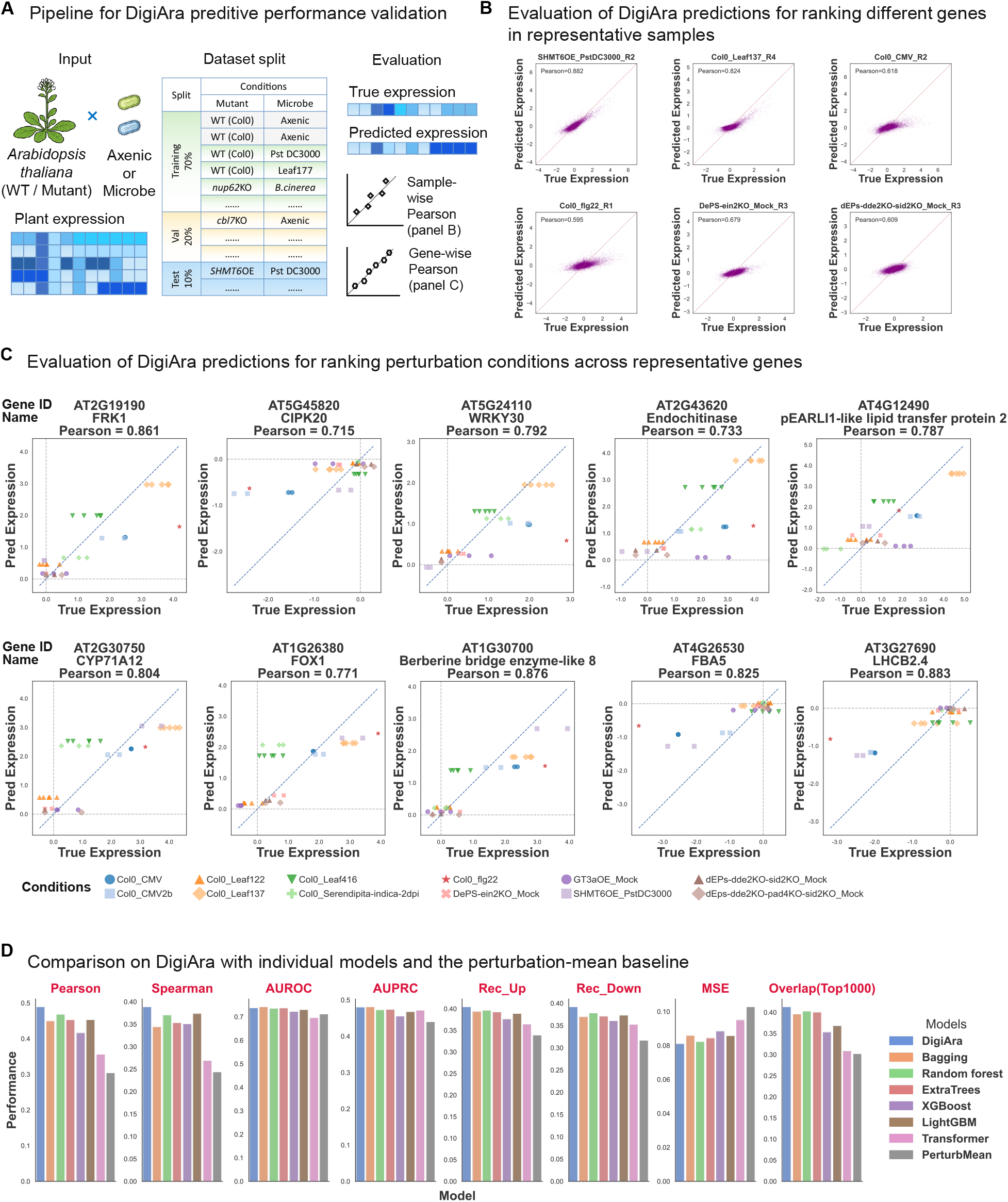
DigiAra accurately predicts *Arabidopsis* transcriptional responses to unobserved genetic perturbations and microbial infection conditions. (A) DigiAra was evaluated on genetic, microbial, and combined genetic–microbial perturbation conditions held out from model training. Conditions were partitioned at the condition level into training, validation, and test sets at a 7:2:1 ratio, and test-set performance was assessed sample-wise and gene-wise by comparing predicted and observed expression. (Caption continued on the next page *↓*.) (Caption continued from the previous page *↑*.) (B) Across held-out test samples, the DigiAra achieved a mean sample-wise Pearson correlation of 0.49 when predicting genome-wide transcriptional responses to genetic and microbial perturbations. Six representative test samples were selected to cover distinct perturbation classes, including a combined genetic–microbial perturbation (*SHMT6* overexpression with *Pst* DC3000), microbial or immune-elicitor perturbations in Col-0 (Leaf137, CMV, and flg22), and genetic perturbations under mock conditions (*ein2* knockout and *dde2 sid2* double knockout). Each point represents one gene; the Pearson correlation for each sample is indicated above the corresponding plot, and the dashed diagonal denotes the identity line (*y* = *x*). (C) Across all genes, the DigiAra achieved a mean gene-wise Pearson correlation of 0.32 between predicted and observed expression across held-out test samples. Ten representative genes were selected to span four functional layers: immune and stress signaling or transcriptional regulation (*FRK1, CIPK20*, and *WRKY30*); defense effectors (Endochitinase and pEARLI1-like lipid transfer protein 2); specialized defense metabolism and redox-associated processes (*CYP71A12, FOX1*, and *ATBBE8* (Berberine bridge enzyme-like 8)); and core carbon metabolism or photosynthesis (*FBA5* and *LHCB2*.*4*). These genes also encompass diverse condition-dependent expression patterns, including strong induction, moderate responses, and repression. Their gene-wise Pearson correlations ranged from 0.715 to 0.883. Each point represents one test sample; identical colors and marker shapes denote samples from the same genetic–microbial perturbation condition. The diagonal dashed line denotes *y* = *x*, and the horizontal and vertical dashed lines indicate zero expression. (D) Across eight complementary performance metrics, the DigiAra showed the best overall performance among six individual deep and shallow models and the perturbation-mean baseline. The evaluated metrics included Pearson and Spearman correlations, AUROC, AUPRC, recall of up- and down-regulated genes, mean squared error, and top-1,000 gene overlap.

At the sample level, DigiAra achieve a mean Pearson correlation of 0.49 between predicted and observed genome-wide expression profiles across all held-out test samples (Fig. 2B; Fig. S1). Figure 2B presents six representative predictions from the full test set, each comprising approximately 27,448 plant genes and collectively spanning distinct perturbation classes. These examples include the combined genetic–microbial perturbation *SHMT6OE_PstDC3000_R2* (Pearson correlation = 0.882), which involves an *SHMT6*-overexpression background infected with *Pst* DC3000, a widely used bacterial model pathogen for studying plant immunity in *Arabidopsis* [19]. The microbial and immune-elicitor examples in wild-type Col-0 included *Col0_Leaf137_R4* (Pearson correlation = 0.824), *Col0_CMV_R2* (Pearson correlation = 0.618), and *Col0_flg22_R1* (Pearson correlation = 0.595). The CMV condition represents infection with cucumber mosaic virus, a plant RNA virus with broad effects on host growth and physiology [20], whereas flg22 activates pattern-triggered immune signaling. The genetic perturbations under mock conditions included *DePS-ein2KO_Mock_R3* (Pearson correlation = 0.679) and *dEPs-dde2KO-sid2KO_Mock_R3* (Pearson correlation = 0.609), representing perturbations of ethylene signaling and jasmonate/salicylic acid signaling, respectively. Across these representative examples, the hybrid model outperformed conventional machine-learning approaches (Fig. S1; Fig. S2A). The overall test-set performance and representative predictions demonstrate that DigiAra can reconstruct transcriptome-wide expression states across diverse genetic, microbial, and combined genetic–microbial perturbation conditions held out from model training.

At the gene level, DigiAra achieve a mean gene-wise Pearson correlation of 0.32 across all genes in the held-out test set (Fig. S3B), representing the highest overall gene-wise performance among the evaluated models (Table S1). Ten representative genes were selected to span four functional categories and diverse condition-dependent expression patterns. The immune and stress signaling or transcriptional regulation group included *FRK1* (*AT2G19190*; Pearson correlation = 0.861), *CIPK20* (*AT5G45820*; Pearson correlation = 0.715), and *WRKY30* (*AT5G24110*; Pearson correlation = 0.792). *FRK1* is a well-established marker of flagellin-triggered immune signaling [21], whereas *WRKY30* is a stress-responsive transcription factor associated with defense-related gene regulation and resistance to cucumber mosaic virus in *Arabidopsis* [22]. Representative defense effectors included the endochitinase-encoding gene *AT2G43620* (Pearson correlation = 0.733) and *AT4G12490*, which encodes a pEARLI1-like lipid transfer protein 2 (Pearson correlation = 0.787). Genes representing specialized defense metabolism and redox-associated processes included *CYP71A12* (*AT2G30750*; Pearson correlation = 0.804), *FOX1* (*AT1G26380*; Pearson correlation = 0.771), and *AtBBE8* (*AT1G30700*; Pearson correlation = 0.876). *CYP71A12* contributes to pathogen-responsive, tryptophan-derived metabolism and immune defense [23], whereas *FOX1* and *AtBBE8* encode members of the berberine bridge enzyme-like oxidoreductase family. Finally, the core metabolism and photosynthesis group included *FBA5* (*AT4G26530*; Pearson correlation = 0.825), which encodes a fructose-bisphosphate aldolase involved in central carbon metabolism, and *LHCB2*.*4* (*AT3G27690*; Pearson correlation = 0.883), which encodes a component of the photosystem II light-harvesting complex. Collectively, these genes encompass diverse condition-dependent expression patterns, including strong induction, moderate responses, and low or repressed expression states. The agreement between predicted and observed values indicates that DigiAra preserves the relative expression patterns of individual genes across perturbation conditions, enabling conditions associated with higher or lower expression levels to be distinguished. Thus, DigiAra recovers gene-specific expression dynamics across held-out conditions for genes involved in signaling, transcriptional regulation, defense execution, specialized defense metabolism, central carbon metabolism, and photosynthesis. For direct comparison, the sample-wise and gene-wise predictions produced by random forest, the best-performing comparator model, are provided in Fig. S2 and Table S1, respectively.

We benchmark DigiAra against six conventional machine-learning models and a perturbation-mean baseline using eight complementary performance metrics: Pearson and Spearman correlations, AUROC, AUPRC, recall of up- and down-regulated genes, mean squared error, and top-1,000 gene overlap (Fig. 2D; Table S1). DigiAra achieve the best overall performance across this suite of metrics. To complement these global benchmarks, we further evaluate the reliability of DigiAra predictions at the individual-gene level (Appendix A.7). This analysis identifies 1,403 high-confidence genes whose predicted responses are strongly correlated with experimental measurements and that are enriched in defense, stress-response, and signaling functions, further supporting DigiAra’s ability to capture transcriptional programs relevant to plant immunity (Fig. S3). These results support the use of DigiAra as the predictive foundation for subsequent counterfactual simulation, scoring and screening.

### DigiAra’s predictions recapitulate the general non-self response immune program and are linked to observed bacterial load during infection

Beyond the accuracy of transcriptional response prediction, we further ask whether DigiAra’s predicted transcriptional profiles recapitulate established *Arabidopsis* biological programs associated with microbial infection (Fig. 3A).

**Figure 3:**
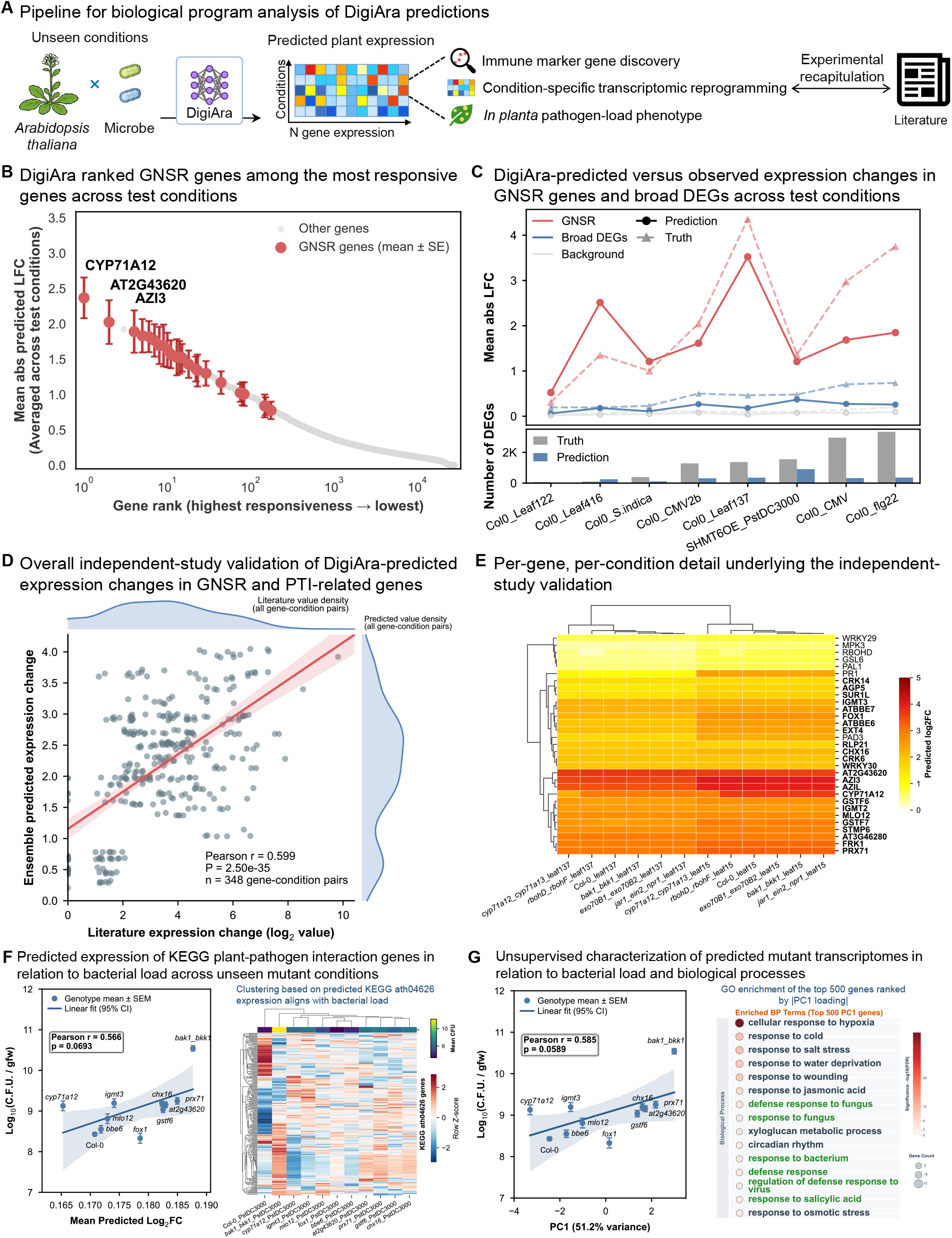
DigiAra’s predictions recapitulate the general non-self response immune program and are linked to observed bacterial load during infection. (Caption continued on the next page *↓*.) (Caption continued from the previous page*↑*.) (A) DigiAra predicts transcriptomes for dataset-unseen plant–microbe conditions. The resulting predictions are evaluated through recovery of immune marker programs, analysis of condition-specific transcriptional reprogramming, and comparison of model-inferred immune activity with experimentally measured bacterial-load phenotypes. (B)DigiAra ranks the previously defined general non-self response (GNSR) genes [24] among the most responsive genes across test conditions. GNSR genes are shown in red as mean *±* SE across conditions, whereas all other genes are shown in gray; selected highly responsive GNSR genes are labeled. (C) Across microbial perturbation test conditions, DigiAra recovers elevated GNSR responsiveness relative to a broad set of differentially expressed genes (DEGs) and background genes, and recapitulates the positive relationship between GNSR responsiveness and the number of DEGs (nDEGs). Solid lines with circles represent DigiAra predictions, whereas dashed lines with triangles represent experimentally observed values; the lower panel compares predicted and observed nDEGs. (D) In dataset-unseen mutant–microbe conditions, DigiAra-predicted expression changes for GNSR and selected pattern-triggered immunity (PTI) marker genes correlate with expression changes measured in an independent study [25] across 348 gene–condition pairs (Pearson *r* = 0.599, *P* = 2.50 *×* 10^*−*35^). Each point represents one marker gene under one mutant–bacterium condition. The line and shaded band indicate the linear fit and 95% confidence interval, respectively, and the marginal plots show the distributions of experimentally measured and predicted expression changes. (E) The corresponding gene-by-condition heatmap provides a detailed view of the DigiAra predictions evaluated in (D), showing condition-specific expression patterns for GNSR genes (bold) and selected PTI marker genes (regular weight) across the same dataset-unseen mutant–microbe conditions. (F) Across *Arabidopsis* GNSR mutants, known immunocompromised *bak1 bkk1* mutant and the wild-type reference Col-0 challenged with *Pst* DC3000, the mean predicted log_2_ fold change of genes in the KEGG plant–pathogen interaction pathway (ath04626) shows a positive trend with experimentally measured bacterial load for the same plant genetic background [24] (Pearson *r* = 0.566, *P* = 0.0693). The heatmap shows row-standardized predicted expression of ath04626 genes, with mutants clustered according to the similarity of their predicted expression profiles. The upper annotation indicates mean bacterial load. The resulting expression-based clusters largely group mutants with similar bacterial loads, suggesting concordance between the predicted pathway-level expression and the observed phenotype. (G) For the same plant genetic background and bacterial-load measurements used in (F), PC1 derived from the complete DigiAra-predicted transcriptomes shows a positive trend with observed bacterial load (Pearson *r* = 0.585, *P* = 0.0589). PC1 explains 51.2% of the predicted transcriptional variance. Gene Ontology enrichment analysis of the 500 genes with the largest absolute PC1 loadings identifies biological processes related to defense, immunity, and stress responses. In the scatter plots in (F) and (G), each point represents one plant genetic background, with observed bacterial load shown as mean *±* SEM; the lines and shaded bands indicate linear fits and 95% confidence intervals.

We focus on the general non-self response (GNSR), a 24-gene program whose response amplitude tracks the extent of broader transcriptional reprogramming across bacterial perturbations and serves as a biological reference [24]. Within the held-out test set, DigiAra predicts that GNSR genes rank near the top of the genome-wide distribution based on their mean absolute predicted log_2_ fold changes across test conditions (Fig. 3B). Among these genes, *CYP71A12*, the endochitinase-encoding gene *AT2G43620*, and *AZI3* are among the most strongly predicted responders. Across microbial perturbation conditions, DigiAra-predicted GNSR response amplitudes are consistently greater than those of a broader DEG set and background genes and show an overall positive relationship with the total number of differentially expressed genes (nDEGs) (Fig. 3C; Fig. S4A). Although DigiAra generally underestimates absolute response amplitudes and nDEG counts, it nevertheless recovers the condition-dependent trend in GNSR responsiveness and its relationship with the extent of transcriptional reprogramming.

The immune-response structure recovered in the held-out test set extends to an independent mutant–bacterium dataset. An independent study quantifies GNSR and pattern-triggered immunity (PTI) marker responses across diverse *Arabidopsis* mutant–bacterium combinations [25]. Because these combinations are absent from the DigiAra training dataset, they provide an external test of model generalizability. Across 348 marker gene–condition pairs, DigiAra-predicted expression changes correlate with experimentally measured values (Pearson *r* = 0.599, *P* = 2.50 *×* 10^*−*35^; Fig. 3D). The predicted profiles further capture distinct mutation- and bacterium-dependent expression patterns, revealing context-specific immune transcriptional responses across unseen conditions (Fig. 3E).

Experimentally measured *in planta* bacterial loads provide an additional trait-level benchmark [24]. Across nine GNSR mutants (*prx71, chx16, gstf6, bbe6, mlo12, at2g43620, igmt3, fox1*, and *cyp71a12*), the known immunocompromised *bak1 bkk1* double mutant, and the wild-type reference Col-0 challenged with *Pst* DC3000, the mean predicted expression change across genes in the KEGG plant–pathogen interaction pathway (ath04626) shows a positive trend with the observed bacterial load (Pearson *r* = 0.566, *P* = 0.0693; Fig. 3F). To determine whether this trait-related variation can be recovered without prior pathway specification, we apply principal component analysis to the complete genome-wide expression profiles predicted for the same *Pst* DC3000-challenged genotypes. PC1 explains 51.2% of the total predicted transcriptional variance and shows a concordant positive trend with the same bacterial-load measurements (Pearson *r* = 0.585, *P* = 0.0589; Fig. 3G; Fig. S4B; Fig. S4C). The 500 genes with the largest absolute PC1 loadings are enriched for defense-, immune-, and stress-related biological processes, independently recovering the functional programs highlighted by the pathway-informed analysis. These analyses show that DigiAra extends beyond expression reconstruction to recover immune-related patterns across multiple scales, from individual marker genes and condition-specific response programs to genome-wide transcriptional states that show positive trends with *in planta* pathogen burden. This multiscale biological concordance supports the use of DigiAra for biologically informed virtual screening and genotype prioritization.

### DigiAra prioritizes *Arabidopsis* mutants predicted to enhance immunity while minimizing growth suppression

Having established that DigiAra accurately predicts *Arabidopsis* transcriptional responses under held-out conditions and recapitulates immune-response programs across independent biological contexts, we next apply the model prospectively to genome-wide mutant design. Specifically, we seek to identify *Arabidopsis* mutants predicted to enhance immunity against microbial infection while minimizing the suppression of growth-related processes.

To achieve this objective, we use the DigiAra S3 pipeline (Appendices A.5 and A.6). We simulate single-gene knockouts of all 27,448 *Arabidopsis* genes under infection with *Pst* DC3000. For each candidate, we compare the predicted KO–*Pst* DC3000 expression profile with the ensemble-predicted WT–*Pst* DC3000 profile, yielding a gene-level response-difference vector that estimates the knockout-dependent transcriptional effect relative to the shared infection background (Fig. 4A). We then summarize the predicted knockout effects using a custom KEGG pathway-based evaluation module comprising six immunity-related pathways, one plant hormone signaling pathway, and 11 growth-associated pathways (Table S2). Candidate screening proceeds in two steps. First, we retain knockouts with a positive mean immunity Signed-RMS score, narrowing the genome-wide search space from 27,448 genes to 27 candidates (0.098%). Second, we rank these 27 candidates using an immune–growth balance score that favors stronger predicted immune responses while penalizing disruption of growth-associated pathways. The hormone signaling scores are retained to provide biological context but are not used for candidate selection or ranking.

**Figure 4:**
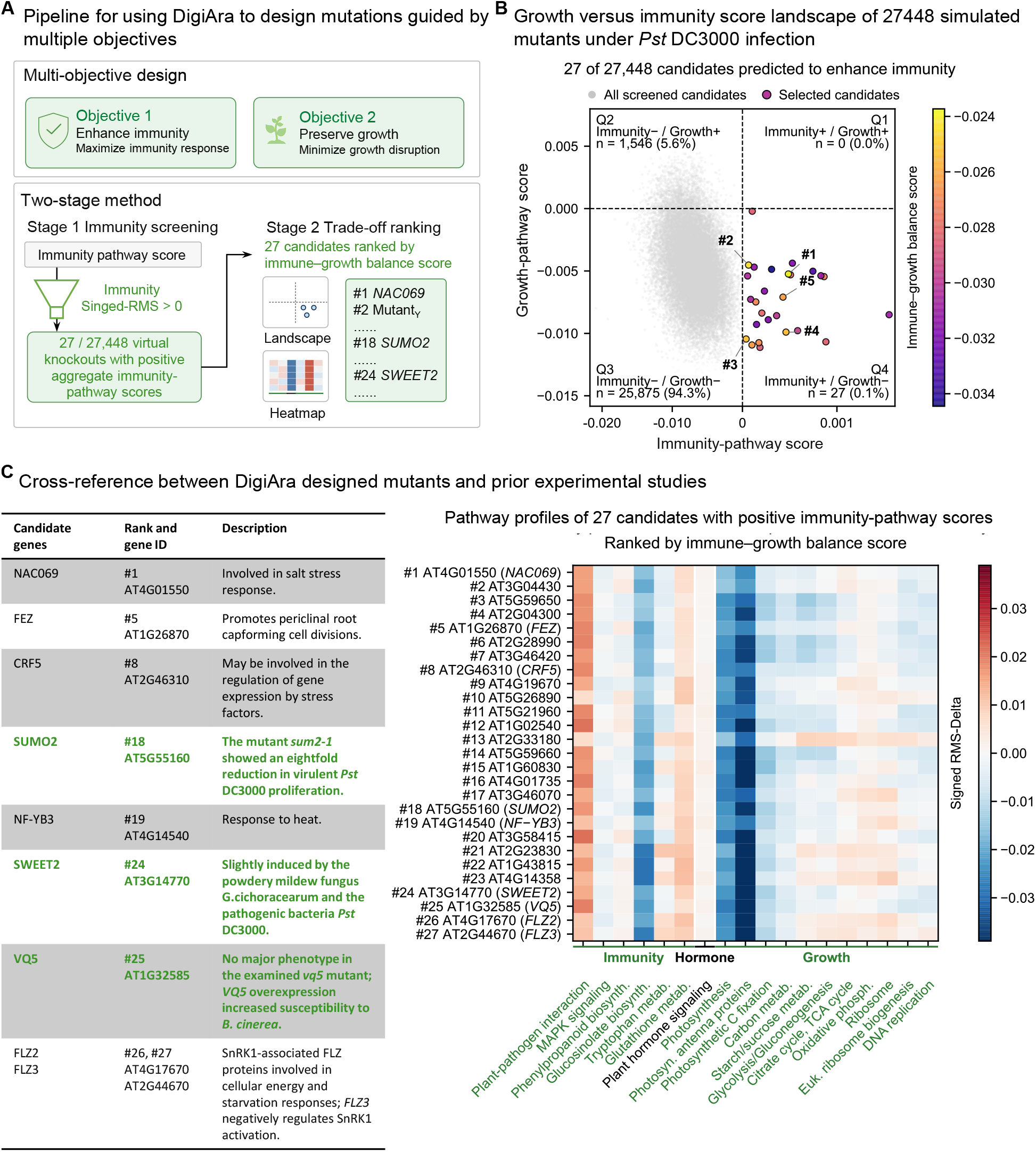
DigiAra prioritizes *Arabidopsis* mutants predicted to enhance immunity while minimizing growth suppression. (A) DigiAra uses a multi-objective, two-stage strategy to design and prioritize mutant candidates, seeking enhanced immune-associated responses while limiting growth-pathway perturbation. For each of 27,448 knockouts, the predicted KO–*Pst* DC3000 transcriptome was compared with the predicted WT–*Pst* DC3000 transcriptome. Knockout effects were summarized as RMS-Delta and Signed-RMS across six immunity-associated, one plant hormone signaling, and 11 growth-associated pathways. In stage 1, immunity feasibility screening retained candidates with positive mean immunity Signed-RMS. In stage 2, immunity–growth trade-off ranking prioritized the retained candidates using an immunity–growth balance score. Plant hormone signaling was retained as contextual information but excluded from candidate selection and ranking. (Caption continued on the next page *↓*.) (Caption continued from the previous page *↑*.) (B) Genome-wide simulation identified 27 immunity-positive knockout candidates, which were ranked by balancing predicted immune enhancement against the magnitude of growth-pathway perturbation. The signed immunity–growth landscape shows all 27,448 knockouts, with grey points representing all screened candidates and coloured points with black outlines representing the 27 selected candidates. Candidate colour indicates the immune–growth balance score, and the five highest-ranked candidates are labeled. Dashed lines mark zero, and quadrant annotations report candidate counts and percentages. The negative and positive domains of the immunity axis are displayed at equal widths, whereas the growth axis remains linear. (C) Candidates with curated gene symbols and reported functions include *NAC069* [26], *FEZ* [27], *CRF5* [28], *SUMO2* [29], *NF-YB3* [30], *SWEET2* [31, 32], *VQ5* [33], *FLZ2* [34], and *FLZ3*/*DUF581-9* [35]; their reported functions and sources are summarized at left. The heatmap shows Signed-RMS profiles for all 27 candidates, ordered by immune–growth balance score, across six Immunity, one Hormone, and 11 Growth pathways. Plant–pathogen interaction responses are predominantly positive, whereas glucosinolate biosynthesis, photosynthesis, and photosynthesis–antenna protein responses are predominantly negative. Red and blue indicate positive and negative Signed-RMS values, respectively.

The signed immunity–growth landscape reveals a strongly asymmetric distribution of predicted knockout effects (Fig. 4B). Most knockouts occupy the immunity-negative/growth-negative Signed-RMS quadrant (25,875; 94.3%), whereas 1,546 candidates (5.6%) exhibit negative immunity and positive growth-associated Signed-RMS scores. No candidate exhibits positive Signed-RMS scores for both immunity and growth. Instead, all 27 retained candidates occupy the immunity-positive/growth-negative quadrant. The growth Signed-RMS score is shown to characterize predicted growth-associated responses but is not used as a selection threshold. Among the 27 immunity-positive candidates, the immune–growth balance score favors those with smaller overall perturbations of growth-related pathways. Thus, the screen does not identify knockouts predicted to enhance both immune- and growth-associated transcription; rather, it prioritizes candidates predicted to enhance immunity while limiting suppression of growth.

The pathway-resolved heatmap shows that the positive aggregate immunity scores arise from distinct combinations of pathway responses rather than uniform activation across all immunity-associated pathways (Fig. 4C). Plant–pathogen interaction responses are predominantly positive, whereas glucosinolate biosynthesis, photosynthesis, and photosynthesis– antenna protein responses are predominantly negative. The remaining immunity-related, plant hormone signaling, and growth-associated pathways exhibit heterogeneous responses across the selected candidates, indicating that the predicted profiles retain candidate-specific pathway variation. Previous studies provide functional context for nine of the 27 candidates: *NAC069* (*AT4G01550*) [26], *FEZ* (*AT1G26870*) [27], *CRF5* (*AT2G46310*) [28], *SUMO2* (*AT5G55160*) [29], *NF-YB3* (*AT4G14540*) [30], *SWEET2* (*AT3G14770*) [31, 32], *VQ5* (*AT1G32585*) [33], *FLZ2* (*AT4G17670*) [34], and *FLZ3*/*DUF581-9* (*AT2G44670*) [35]. Notably, *SUMO2* has direct genetic evidence linking its immunity–growth traits considered in our screen: this mutant was reported to support approximately eightfold lower proliferation of virulent *Pst* DC3000 than wild-type plants while showing no obvious growth reduction under the tested conditions [29] (Fig. S5), which is consistent with the placement of *SUMO2* among the 27 candidates with positive aggregate immunity scores and relatively limited predicted disruption of growth-associated pathways.

Among the remaining candidates, *SWEET2* also has direct genetic evidence connecting it to pathogen responses across multiple pathosystems. During infection of *Arabidopsis* leaves by *Xanthomonas campestris* pv. *campestris*, loss-of-function *sweet2* mutants showed reduced apoplastic glucose accumulation, lower bacterial populations, and decreased disease severity [36]. In contrast, three independent *sweet2* insertion mutants were more susceptible to the root-infecting oomycete *Pythium irregulare* [37], whereas *Arabidopsis sweet2* mutants showed enhanced resistance to the clubroot pathogen textitPlasmodiophora brassicae [32]. These contrasting traits connect *SWEET2* to pathogen responses while showing that the direction of its effect differs among pathosystems. Three additional late-ranked candidates provide further functional context, although the available evidence differs in strength. *VQ5* (*AT1G32585*) has the clearest pathogen-related evidence [33]. In a family-wide analysis, a *vq5* insertion mutant showed no major alteration in growth, development, or resistance to virulent *Pseudomonas syringae* or *Botrytis cinerea*, whereas *VQ5*-overexpressing plants developed increased chlorosis following *B. cinerea* infection, consistent with enhanced susceptibility. The overexpression trait is directionally compatible with the predicted immune benefit of *VQ5* loss, but it does not constitute direct validation of the knockout prediction. The absence of a pronounced trait in the previously examined *vq5* mutant may reflect functional redundancy, assay sensitivity, or pathosystem-dependent effects. The remaining two candidates, *FLZ2* (*AT4G17670*) and *FLZ3*/*DUF581-9* (*AT2G44670*), belong to the FCS-like zinc-finger/DUF581 family associated with SnRK1 energy signaling. *FLZ2* interacts with SnRK1 subunits and is regulated by cellular energy status, whereas *FLZ3* directly restricts SnRK1 activation [34, 38, 35]. Loss of *FLZ3* enhanced a dark-induced SnRK1 transcriptional response without improving survival after prolonged darkness, while its overexpression impaired recovery from starvation. Their joint selection therefore points more strongly to an energy-homeostasis and growth–stress regulatory axis than to previously established pathogen-resistance traits.

## Discussion

DigiAra establishes a computational framework for designing genetic mutations associated with desired traits in *Arabidopsis thaliana*. Through its simulation–scoring–screening (S3) pipeline, DigiAra integrates genetic perturbations, microbial conditions, and background transcriptional states to predict genome-wide expression profiles under unobserved conditions, and translate these predictions into trait-oriented candidate rankings. The predicted profiles consistently agree with measured expression at both the sample and gene levels, and recover established immune-response programs. Specifically, DigiAra recapitulates the general non-self response (GNSR), and in an independent dataset, the predicted pattern-triggered immunity pathway scores further correlate with experimentally measured bacterial loads.

Further genome-wide simulation with DigiAra under *Pst* DC3000 infection identifies 27 gene knockouts with positive aggregate immunity scores and ranks them according to their predicted disruption of growth-associated pathways. 9 candidates have reported functions related to immunity, stress responses, development, or energy regulation, among which *SUMO2* is directly supported by previously reported experimental evidence. Nevertheless, the prioritized candidates should currently be regarded as testable hypotheses rather than experimentally validated genes.

## Acknowledgements

This work was supported in part by the University Development Fund–Research Start-Up Fund from The Chinese University of Hong Kong, Shenzhen (Grant No. UDF01004259).

## Appendix

### A Methods

#### A.1 RNA-seq data collection and processing

Publicly available bulk RNA-seq data of *Arabidopsis thaliana* were assembled from 26 independent studies, comprising 495 samples, 49 plant genetic background, 45 microbial challenges and 137 distinct experimental conditions. The microbial challenges included bacteria, fungi, viruses and peptide elicitors.

All sequencing data were processed using a uniform workflow. The TAIR10 *A. thaliana* cDNA reference was obtained from Ensembl Plants release 62 (https://ftp.ebi.ac.uk/ensemblgenomes/pub/release-62/plants/fasta/arabidopsis_thaliana/cdna/) and indexed using Salmon v1.10.3 with a *k*-mer size of 31 [18]. Reads from each sample were quantified against this index, and the NumReads field was retained as the Salmon-estimated transcript count.

Transcript identifiers were linked to the *A. thaliana* UniProt reference proteome (UP000006548; https://ftp.uniprot.org/pub/databases/uniprot/current_release/knowledgebase/reference_proteomes/Eukaryota/UP000006548/UP000006548_3702.fasta.gz) to align the RNA-seq measurements with the protein embeddings used by the prediction models. For plant gene *i* in sample *s*, estimated counts from all associated transcripts were summed:

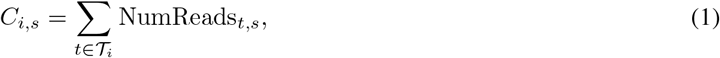

where *T*_*i*_ denotes the set of transcripts linked to the corresponding UniProt accession. The aggregated counts were converted to Counts Per Million (CPM) and transformed using the natural-logarithm `log1p` function:]

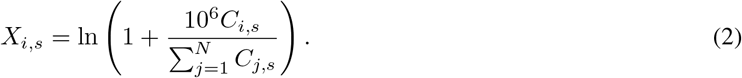

Here, *X*_*i*,*s*_ denotes the normalized RNA-seq expression value of plant gene *i* in sample *s*. The complete expression profile of each sample was represented as a plant transcriptomic state vector,

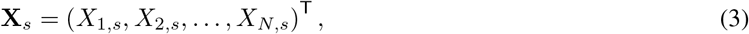

spanning *N* = 27,448 plant genes. A fixed gene order linked to the UniProt reference proteome was maintained across the expression matrix and plant protein embeddings.

#### A.2 Feature representation for plant–microbe interactions

The model inputs integrated the experimentally determined plant transcriptomic state with sequence-derived representations of plant proteins and microbial challenges.

##### A.2.1 Construction of model inputs

For each RNA-seq study *p*, transcriptomic states from the corresponding wild-type (WT) axenic controls were averaged to define the baseline plant state:

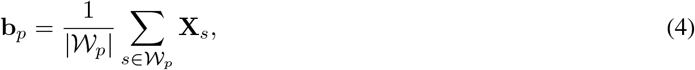

where *W*_*p*_ denotes the set of WT–axenic control samples in study *p*. For experimental condition *c*, the plant-state input **S**_*p*,*c*_ was initialized from **b**_*p*_ and modified when the condition contained a known genetic perturbation.

For each targeted gene, knockout was encoded by replacing the corresponding expression value with *−* ln(1 + 10^6^), whereas overexpression was encoded using + ln(1 + 10^6^). These fixed extrema encoded the intervention and were not interpreted as measured expression values. The remaining genes retained their values from the corresponding baseline state. Multiple genetic perturbations were represented by replacing all targeted coordinates simultaneously. Knockout and overexpression encodings were both included during model training, whereas subsequent counterfactual screening used the knockout encoding.

After construction of the condition-specific plant state, plant sequence information was incorporated using ESM-2 (esm2_t48_15B_UR50D; https://huggingface.co/facebook/esm2_t48_15B_UR50D) [17]. The resulting plant embedding matrix **E**_plant_ *∈* ℝ^*N ×d*^, where *d* = 5,120, contained one sequence-derived representation for each coordinate of the plant transcriptomic state. The plant embeddings were modulated by the corresponding WT or mutant-specific state:

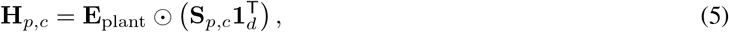

where *⊙* denotes element-wise multiplication and **1**_*d*_ is a *d*-dimensional vector of ones.

For microbial challenge *m*, ESM-2 representations of the corresponding microbial proteins, or of the elicitor peptide where applicable, were mean-pooled to obtain a microbial embedding **e**_*m*_ *∈* ℝ^*d*^. This vector was broadcast across the plant dimension and concatenated with the state-modulated plant representation:

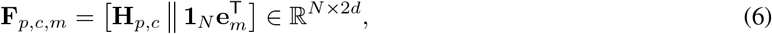

where ∥ denotes feature concatenation and **1**_*N*_ is an *N*-dimensional vector of ones. The Transformer received the complete matrix **F**_*p*,*c*,*m*_, whereas the shallow machine-learning models received its mean across the plant-gene dimension, producing a 2*d*-dimensional input vector.

##### A.2.2 Plant expression changes predicted by the models

The models were trained to predict transcriptome-wide expression changes relative to the corresponding baseline plant state. For sample *s*, the prediction target was defined as

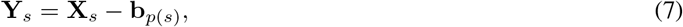

where *p*(*s*) denotes the RNA-seq study from which sample *s* was obtained. Thus, **Y**_*s*_ represented the expression response associated with a genetic perturbation, microbial exposure or their combination, relative to the mean WT– axenic expression profile from the same study. All shallow models and the Transformer produced an *N*-dimensional output containing the predicted expression changes of the 27,448 plant genes in the fixed model order.

#### A.3 Dataset split strategy

137 distinct experimental conditions were stratified into three perturbation categories: mutant-only, microbe-only, and combined mutant–microbe conditions. Within each category, the conditions and all corresponding biological replicates were randomly allocated to the training, validation, and test sets in a 7:2:1 ratio. All wild-type mock samples receiving a sterile control treatment were assigned to the training set.

#### A.4 Transcriptome prediction and construction of the hybrid model

##### A.4.1 Shallow and deep models for transcriptome prediction

Using the input representations and transcriptomic response targets described above, we evaluated a panel of shallow regression models implemented primarily in scikit-learn [39]. The evaluated algorithms included linear regression, Ridge regression, Lasso, Elastic Net, *k*-nearest-neighbour regression, kernel ridge regression, partial least-squares regression, multilayer perceptrons, bagging, random forests and extremely randomized trees (ExtraTrees). XGBoost [40] and LightGBM [41] were additionally evaluated using their respective Python implementations. In parallel, a Transformer architecture [42] was implemented in PyTorch [43] and trained using the matrix-form input described above.

##### A.4.2 Model selection and prediction integration

Candidate models were selected according to the Pearson correlation between predicted and observed transcriptomic responses in the validation set, whereas the test set was reserved for final performance evaluation. Based on validation performance, Bagging, Random Forest, ExtraTrees, XGBoost and LightGBM were retained together with the Transformer. The selected models were trained independently, and their predictions were integrated by an unweighted arithmetic mean:

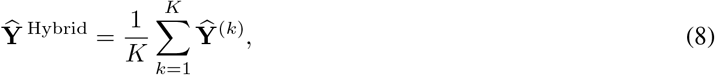

where *K* denotes the number of component models and **Ŷ** ^(*k*)^ is the transcriptomic response predicted by model *k*. The resulting prediction-level combination is referred to throughout as the Hybrid model.

#### A.5 Counterfactual virtual perturbation inference (simulation)

Counterfactual inference was used to estimate how predefined genetic perturbations altered the predicted plant response to a selected microbial challenge. The same knockout encoding used during model construction was applied to the baseline plant state of each eligible reference study. Genome-wide screening and targeted evaluation of selected single- or multi-gene mutants followed the same procedure, with multi-gene perturbations represented by replacing all targeted coordinates simultaneously.

A common plant reference panel was used across microbial challenges. Let *P* _ref_ denote the set of source studies that satisfied the reference criteria. Eligible studies contained a *Pst* DC3000 treatment and a valid corresponding WT–axenic control state. The size of *P*_ref_ was determined by data availability rather than fixed as part of the method.

For variant *v* and microbial challenge *m*, a mutant-specific plant state was constructed independently from each baseline **b**_*p*_, where *p* ∈ *P* _ref_. Each component model generated a transcriptomic response prediction, and the predictions were averaged across both the reference panel and the component models:

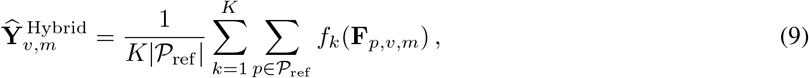

where *f*_*k*_ denotes component model *k*, and **F**_*p*,*v*,*m*_ is the input constructed for variant *v*, microbial challenge *m* and reference study *p*. The same reference panel was used for every microbial challenge, with the microbial embedding varied according to the challenge being evaluated.

A matched WT prediction was generated from the corresponding WT baseline states in the same reference panel and using the same microbial embedding. The predicted effect of the genetic perturbation was then defined as

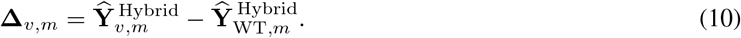

Thus, **Δ**_*v*,*m*_ ∈ ℝ^*N*^ represents the predicted transcriptomic effect of variant *v* relative to matched WT plants exposed to the same microbial challenge. These matched-WT response vectors were used for all subsequent functional analyses and comparisons among candidate mutants.

#### A.6 Biological knowledge-guided evaluation of predicted mutant eesponses (scoring and screening)

##### A.6.1 Focused KEGG pathway panel

Predicted mutant responses were evaluated using a curated panel of 18 Arabidopsis KEGG pathways [44]. The panel contained six immunity pathways, one plant-hormone-signaling pathway, and eleven growth-associated pathways. The immunity pathways represented immune signaling and defense metabolism, whereas the growth pathways represented photosynthesis and carbon assimilation, central carbon and energy metabolism, and biosynthesis and replication. These submodules were used only to organize pathway order and visualization and did not introduce an additional averaging step.

The plant-hormone-signaling pathway was retained as broad biological context because it contains salicylic-acid and other hormone-response branches. It was scored and displayed but was not used for candidate selection, balance-score calculation, or ranking. A pathway was evaluated only when at least three member genes could be mapped to the fixed model gene order. Pathway identifiers, functional assignments, KEGG gene counts, and pathway-specific notes are provided in Table S2. Member genes for each pathway can be retrieved using the corresponding KEGG REST link endpoint.

##### A.6.2 Scoring of predicted pathway changes

For variant *v*, microbial challenge *m*, and gene *j*, the predicted matched-WT response was defined as

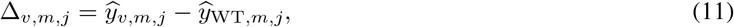

where *ŷ* _*v*,*m*,*j*_ and *ŷ* WT,_*m*,*j*_ denote the predicted challenged-state responses of the mutant and matched WT, respectively. For a KEGG pathway *P* containing *n*_*P*_ mapped genes, the magnitude of the predicted pathway response was quantified using the root mean square (RMS):

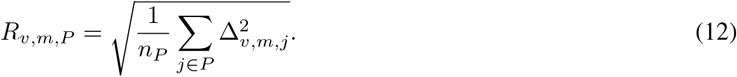

To retain information about response direction, positive and negative components were calculated separately:

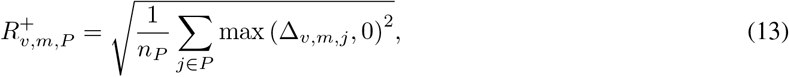

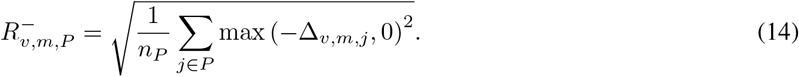

The balance between positive and negative components was summarized as

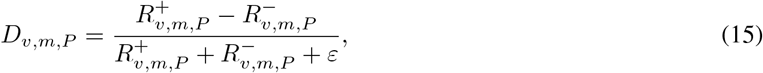

where *ε* = 10^*−*12^ was included for numerical stability. The Signed-RMS was subsequently defined as

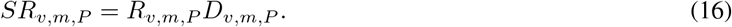

The unsigned RMS therefore quantified the overall magnitude of the predicted pathway change irrespective of sign, whereas the Signed-RMS distinguished responses dominated by increased or decreased expression relative to the matched WT prediction. Each pathway was scored independently. Pathway scores within the immunity and growth groups were then averaged with equal pathway weight.

##### A.6.3 Selection, ranking, and visualization of candidate mutants

Let *P*_*I*_ and *P*_*G*_ denote the six immunity pathways and eleven growth-associated pathways, respectively. The immunity score of variant *v* under microbial challenge *m* was defined as the mean Signed-RMS across the immunity pathways:

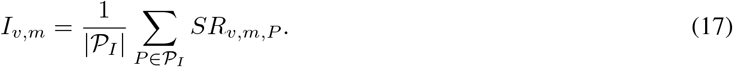

The corresponding growth-disruption score was calculated as the mean unsigned RMS across the growth-associated pathways:

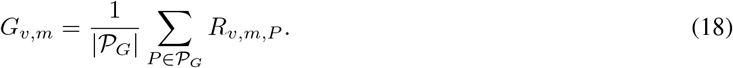

Variants with a positive immunity score were retained as immune-positive candidates:

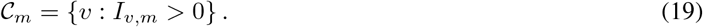

The number of retained candidates was therefore determined by the predicted responses rather than fixed in advance. Within this candidate set, the immune–growth balance score was defined as

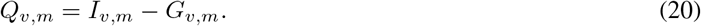

Candidates were ordered by decreasing *Q*_*v*,*m*_, followed, in the event of ties, by decreasing *I*_*v*,*m*_ and then increasing *G*_*v*,*m*_. This ranking favored positive predicted immune responses accompanied by smaller deviations in growth-associated pathways. Because the growth penalty was based on unsigned RMS, substantial growth-associated changes in either direction were treated as departures from growth-associated homeostasis.

For visualization of response direction, a signed growth score was calculated as

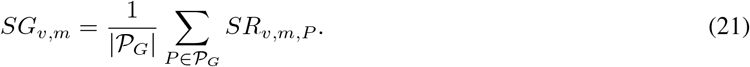

The candidate landscape plotted the immunity score *I*_*v*,*m*_ against the signed growth score *SG*_*v*,*m*_ and used *Q*_*v*,*m*_ as the color scale. Immune-positive candidates were distinguished from the complete screened set, and the highest-ranked candidates were labeled according to the immune–growth balance ranking. For display only, the negative and positive domains of the immunity axis were scaled linearly and shown with equal visual width; distances remained comparable within, but not across, the two sign domains. Pathway heatmaps displayed the Signed-RMS of the 18 retained pathways, with candidate rows ordered by decreasing *Q*_*v*,*m*_.

Candidate mutants were further compared using Pearson correlations and RMS distances between their complete matched-WT response vectors. Genes were ranked by |Δ_*v*,*m*,*j*_| to identify the strongest predicted expression changes associated with each variant–microbe combination.

#### A.7 Statistical evaluation and identification of “highly predictable genes”

To rigorously assess the predictive performance at the single-gene level without relying on the assumption of bivariate normality, we employed a non-parametric empirical bootstrapping strategy. Our goal was to identify a subset of “Highly Predictable Genes” whose dynamic behaviors are captured by the model with both high accuracy and exceptional stability across the dataset.

For each gene, we first calculated the gene-wise Pearson correlation coefficient (*r*) between the predicted and true transcriptomic responses. To decouple the confidence interval (CI) width from the correlation magnitude itself and to mitigate prediction instability driven by sample outliers, we generated *B* = 1000 bootstrap datasets by resampling the prediction-truth pairs with replacement. The 95% Bootstrap Confidence Interval (BCI), denoted as [*BCI*_*low*_, *BCI*_*high*_], was defined by the 2.5th and 97.5th percentiles of this empirical distribution. The Bootstrap CI Width was subsequently computed as *BCI*_*width*_ = *BCI*_*high*_ *− BCI*_*low*_.

A gene was classified as an **“Highly predictable gene”** only if it satisfied the following stringent triple-criteria:

1. **High linear correlation:** Pearson correlation coefficient *r >* 0.7;
2. **High minimum confidence:** Bootstrap CI lower bound *BCI*_*low*_ *>* 0.5;
3. **High stability (narrow variance):** Bootstrap CI width *BCI*_*width*_ *<* 0.4.

These rigorous filters ensure that the “Highly Predictable Genes” represent highly robust biological signals that the model can confidently generalize without being biased by noise.

### B Related work

There is growing interest in developing computational frameworks capable of modeling and predicting system-level cellular responses *in silico*, enabling scalable exploration of large perturbation spaces beyond the limits of conventional experimentation. Representative approaches, including CellOT, scGPT, Geneformer, and CIFM, have modeled cellular state transitions or learned transferable representations of perturbation-responsive transcriptional programs from large-scale transcriptomic data [45, 46, 47, 48, 49]. Together, these advances have established the feasibility of AI-based virtual cells for predicting cellular responses to genetic and chemical perturbations, particularly in mammalian systems.

This progress is beginning to extend into plant biology, where AI-based studies across multiple biological scales are laying important groundwork for the development of generalizable virtual-cell frameworks. Graph-based learning has been used to characterize transcriptional programs associated with complex traits such as drought resistance in major crops [50], while transcriptome-based machine-learning models have enabled the prediction of disease-associated phenotypes, including lesion severity [51]. Single-cell foundation models are also beginning to improve the characterization of cellular heterogeneity in *Arabidopsis* [52]. At the molecular-design level, AI-guided protein engineering is providing new opportunities for rational trait design [53]. More broadly, Ramstein et al. proposed a precision-breeding framework in which sequence-based deep learning predicts functional variant effects and prioritizes candidate variants for selection or genome editing [54]. This framework highlights the potential of computational models to connect functional molecular knowledge with targeted plant improvement.

AI-guided design has also been applied directly to plant immunity. Zhu et al. combined de novo protein design, modular immune-receptor engineering, and *in planta* directed evolution to construct synthetic plant immune receptors capable of recognizing proteins from diverse pathogens [55]. This work provides a compelling demonstration of programmable plant immunity and highlights the potential of integrating computational design with targeted genetic interventions. Collectively, these advances establish a broad foundation for AI-enabled plant research. An important next step is to develop generalizable frameworks capable of predicting how targeted genetic perturbations reshape plant transcriptomic programs across distinct microbial challenges or abiotic stress.

### C Additional results

**Figure S1:**
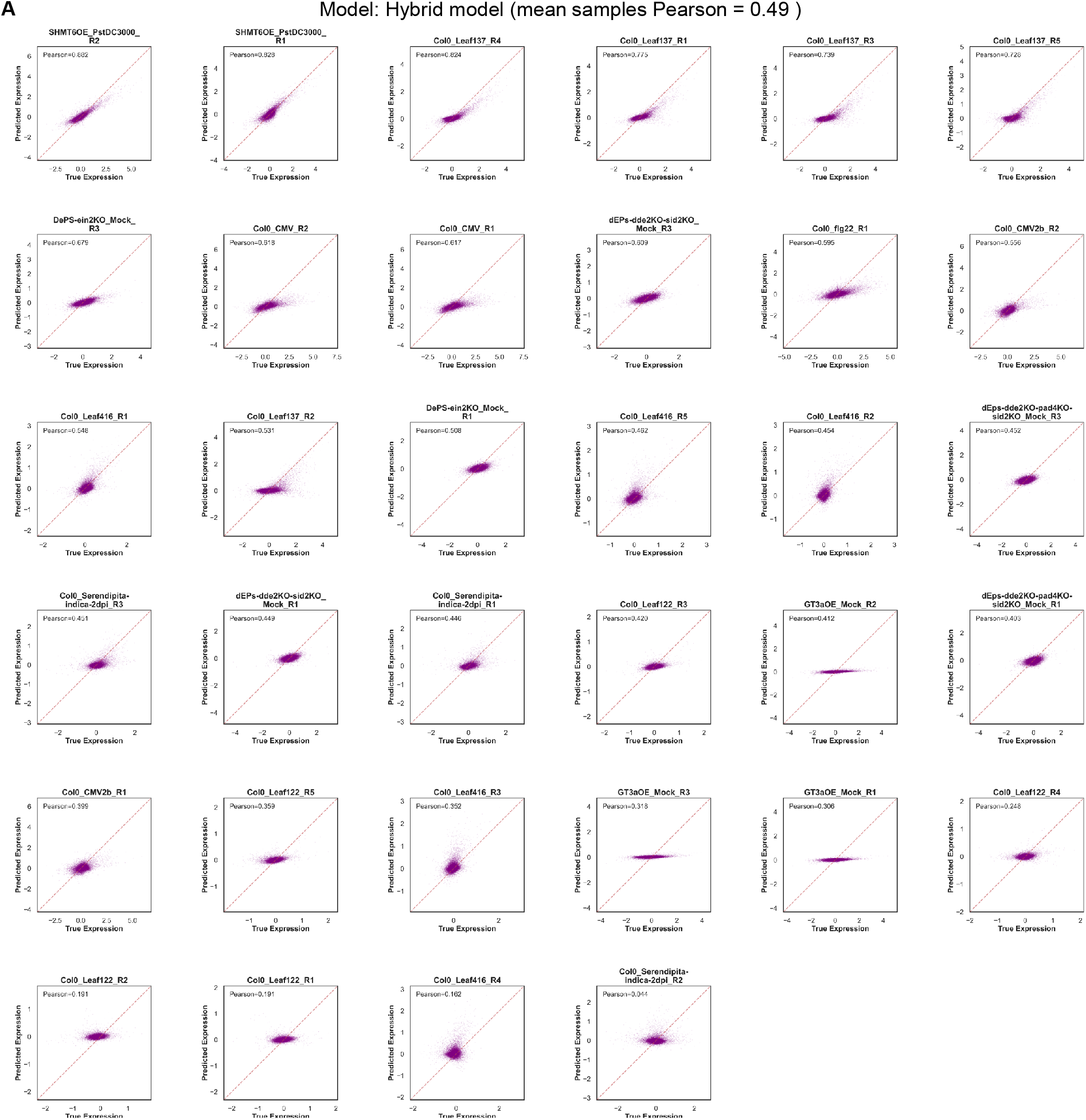
Sample-wise prediction performance of the DigiAra hybrid model across all held-out test samples. (A) Scatterplots compare the genome-wide transcriptional responses predicted by the DigiAra hybrid model with the corresponding experimentally measured responses for each of the 34 held-out test samples spanning genetic and microbial perturbations. Each point represents a gene, and the Pearson correlation coefficient (*r*) between predicted and measured responses is reported in each panel. Samples are ordered by decreasing *r*, and the red dashed lines indicate the identity line (*y* = *x*). The mean sample-wise Pearson correlation across all held-out test samples was 0.49.

**Figure S2:**
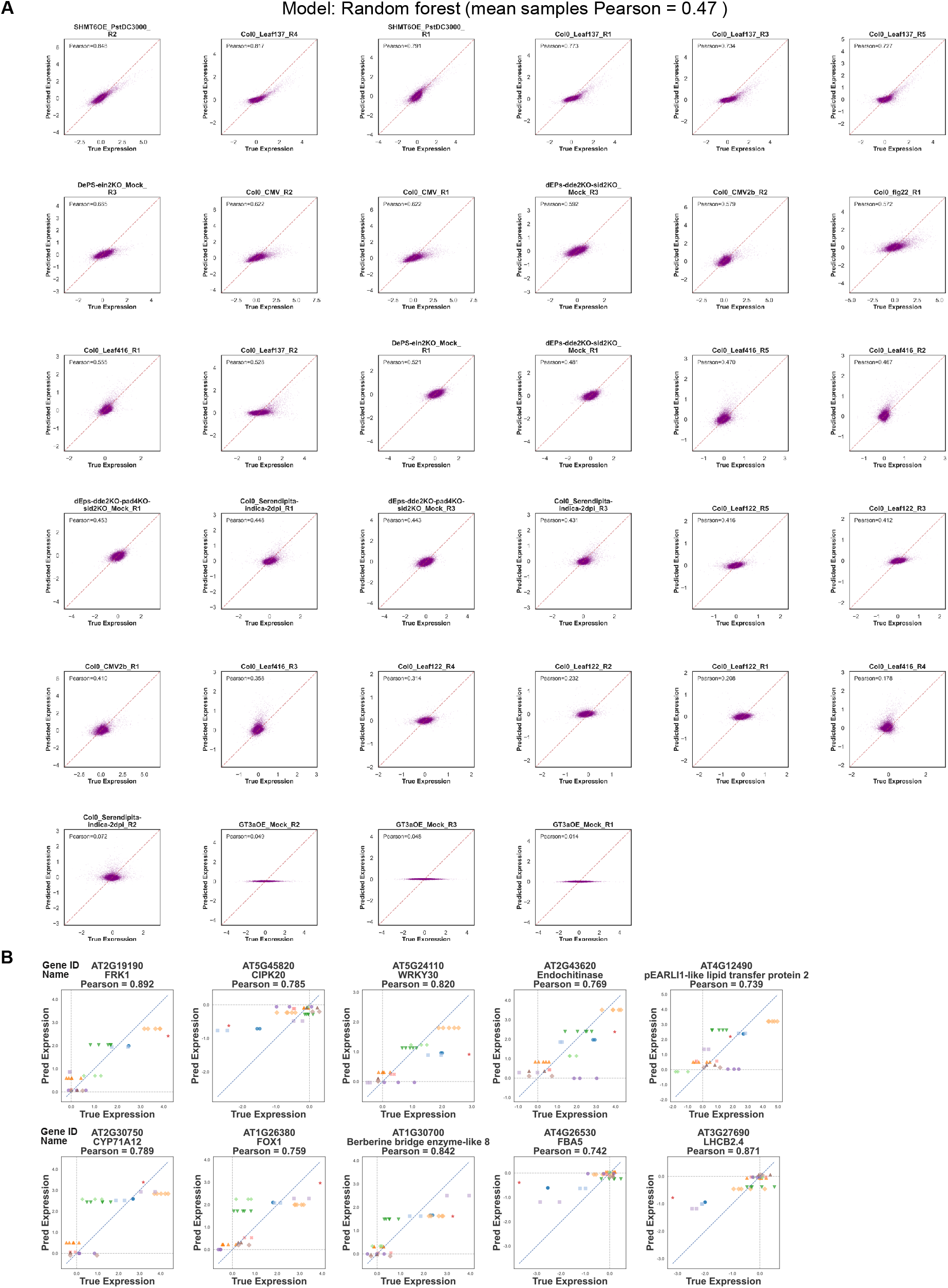
Sample- and gene-wise prediction performance of the random forest model. **(A)** Scatterplots compare random forest-predicted genome-wide transcriptional responses with the corresponding experimentally measured responses for each of the 34 held-out test samples spanning genetic and microbial perturbations. Each point represents a gene, and the Pearson correlation coefficient (*r*) between predicted and measured responses is reported in each panel. Samples are ordered by decreasing *r*, and the red dashed lines indicate the identity line (*y* = *x*). The mean sample-wise Pearson correlation across all held-out samples was 0.47. **(B)** Across all genes, the random forest model achieved a mean gene-wise Pearson correlation of 0.30 between predicted and measured responses across the 34 held-out test samples. Ten representative genes were selected to span four functional layers: immune and stress signaling or transcriptional regulation (*FRK1, CIPK20*, and *WRKY30*); defense effectors (Endochitinase and pEARLI1-like lipid transfer protein 2); specialized defense metabolism and redox-associated processes (*CYP71A12, FOX1*, and *ATBBE8* (Berberine bridge enzyme-like 8)); and core carbon metabolism or photosynthesis (*FBA5* and *LHCB2*.*4*). These genes also encompass diverse condition-dependent expression patterns, including strong induction, moderate responses, and repression. Their gene-wise Pearson correlations ranged from 0.739 to 0.892. Each point represents one held-out test sample; identical colors and marker shapes denote samples from the same genetic–microbial perturbation condition. The diagonal dashed line denotes *y* = *x*, and the horizontal and vertical dashed lines indicate zero response.

**Figure S3:**
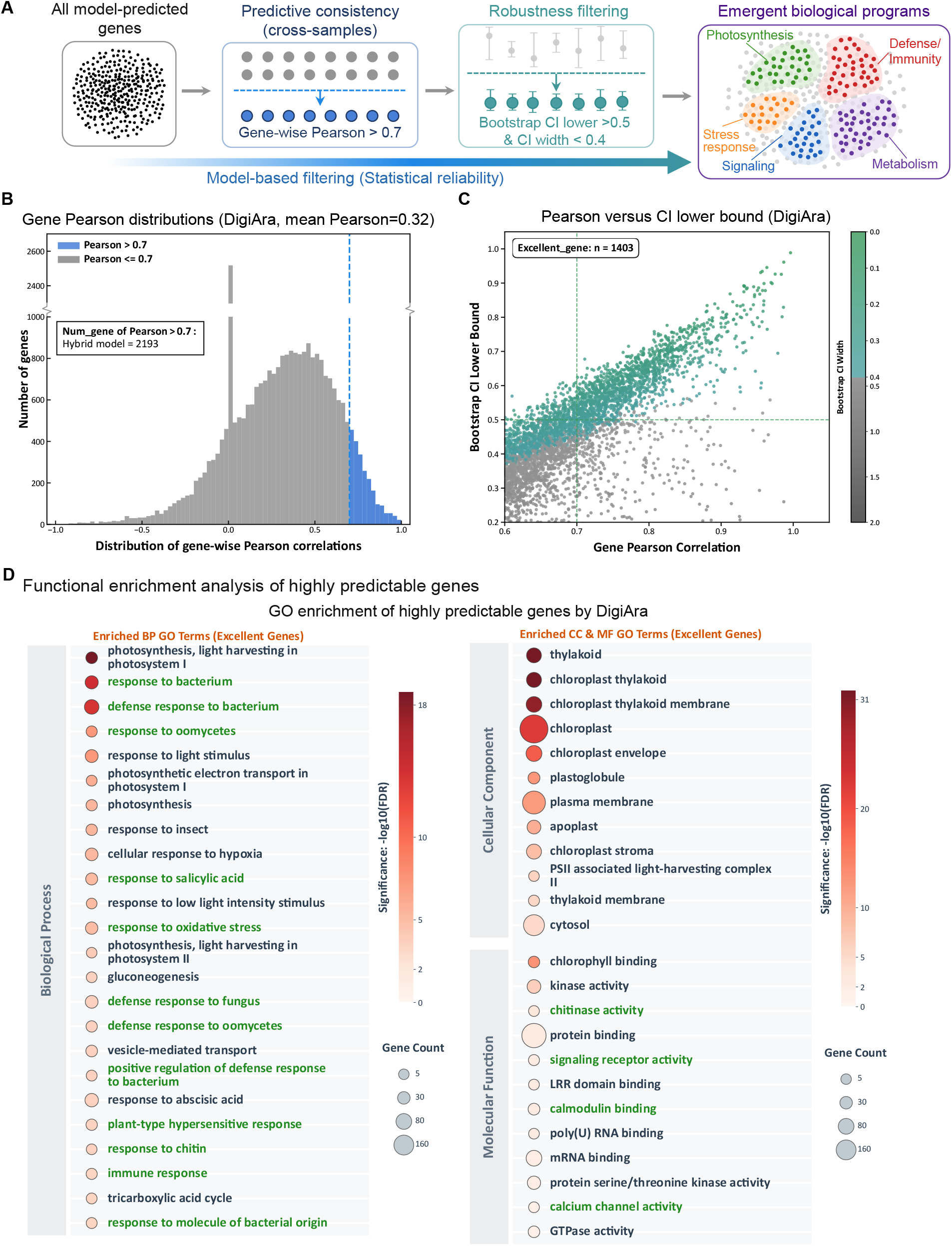
Identification and functional characterization of high-confidence genes predicted by the DigiAra. **(A)** Schematic of the statistical filtering workflow. For each gene, the gene-wise Pearson correlation coefficient (*r*) was calculated between hybrid model-predicted and experimentally measured transcriptional responses across the 34 held-out test samples. Uncertainty was estimated using 1,000 bootstrap resamples of the matched prediction-measurement pairs, with the 95% bootstrap confidence interval (CI) defined by the 2.5th and 97.5th percentiles. Genes were classified as high-confidence genes if they simultaneously satisfied *r >* 0.7, a bootstrap CI lower bound *>* 0.5, and a bootstrap CI width *<* 0.4. **(B)** Distribution of gene-wise Pearson correlations for the DigiAra. The mean gene-wise correlation was 0.32, and 2,193 genes had *r >* 0.7. Genes above this threshold are shown in blue and the remaining genes in gray; the blue dashed line indicates *r* = 0.7. The broken *y*-axis is used to display the full range of gene counts. **(C)** Relationship between the gene-wise Pearson correlation and the lower bound of its 95% bootstrap CI. Each point represents a gene and is colored according to its bootstrap CI width. Dashed lines indicate the correlation (*r* = 0.7) and CI lower-bound (0.5) thresholds. A total of 1,403 genes satisfied all three criteria and were retained as high-confidence genes. **(D)** Gene Ontology enrichment analysis of the 1,403 high-confidence genes. Enriched Biological Process terms are shown on the left, and enriched Cellular Component and Molecular Function terms are shown on the right. Dot size represents the number of genes associated with each term, and color indicates enrichment significance as −log_10_ (FDR). Selected terms related to defense, stress responses, and signaling are highlighted in green.

**Figure S4:**
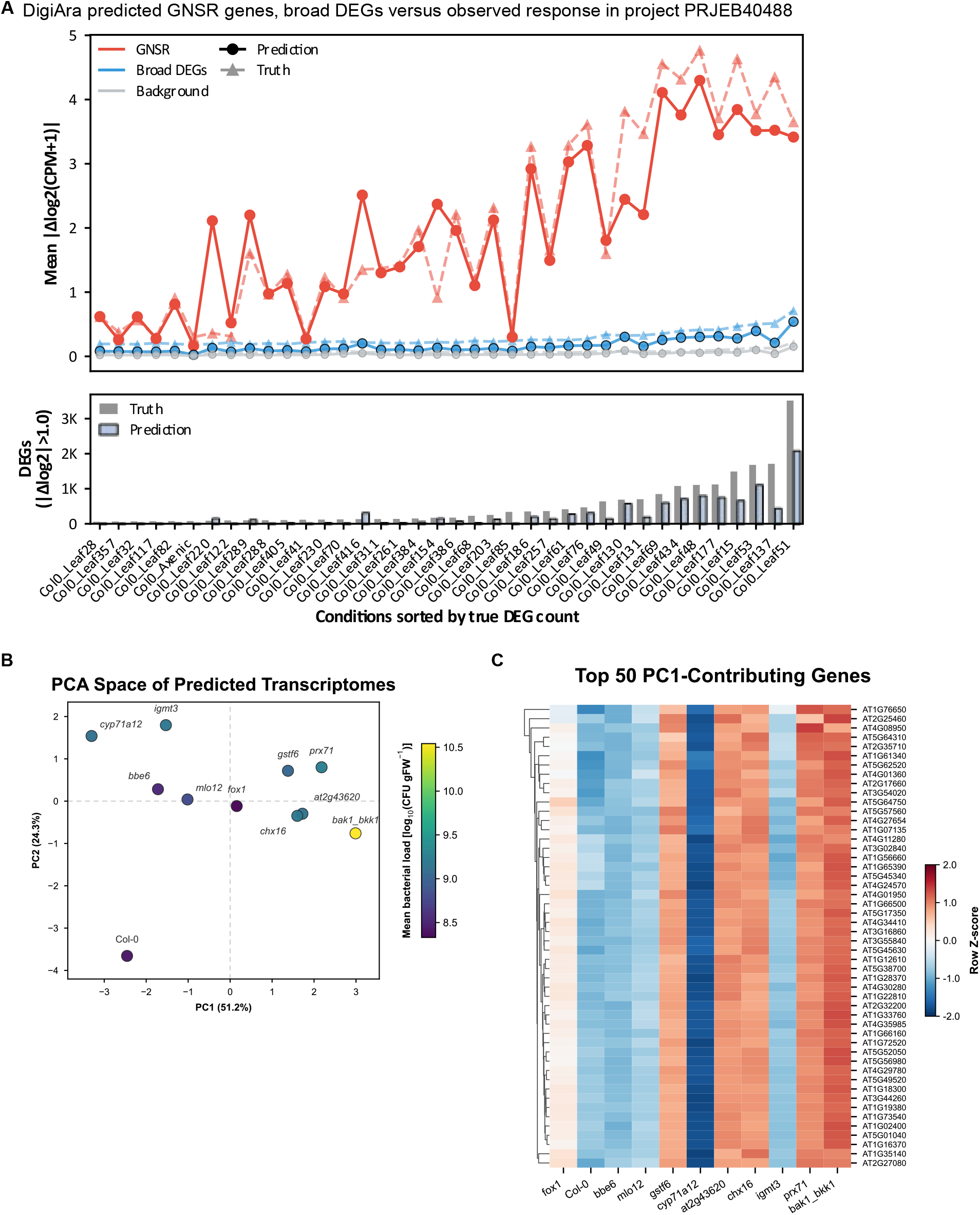
Recovery of GNSR-associated response intensity and colonization-linked transcriptomic structure by the DigiAra hybrid model. **(A)** Condition-level analysis across all 38 conditions in the published PRJEB40488 dataset. For each condition, the upper panel shows the mean absolute transcriptional response, |Δ log_2_(CPM + 1)|, of the 24 general non-self response (GNSR) genes, a broader DEG set, and the remaining background genes. Solid circles represent hybrid model predictions, whereas dashed triangles represent experimentally measured responses. The lower panel compares the measured and predicted numbers of differentially expressed genes (nDEGs), defined by |Δ log_2_| *>* 1. Conditions are ordered by increasing measured nDEG count. The measured data recapitulate the previously reported increase in GNSR amplitude with the extent of transcriptional reprogramming. The hybrid model recovered this condition-dependent trend, although it generally underestimated absolute response amplitudes and nDEG counts. **(B)** Principal component analysis of hybrid model-predicted transcriptomes for 11 Arabidopsis genotypes challenged with *Pst* DC3000. Each point represents one genotype and is colored according to its mean bacterial load, expressed as log_10_(CFU gFW^*−* 1^). PC1 and PC2 explain 51.2% and 24.3% of the total predicted transcriptomic variance, respectively. **(C)** Predicted expression patterns of the 50 genes with the largest absolute PC1 loadings. Values were standardized within each gene and are displayed as row Z-scores. Columns represent genotypes ordered by increasing mean bacterial load, and genes were hierarchically clustered according to their predicted response profiles.

**Figure S5:**
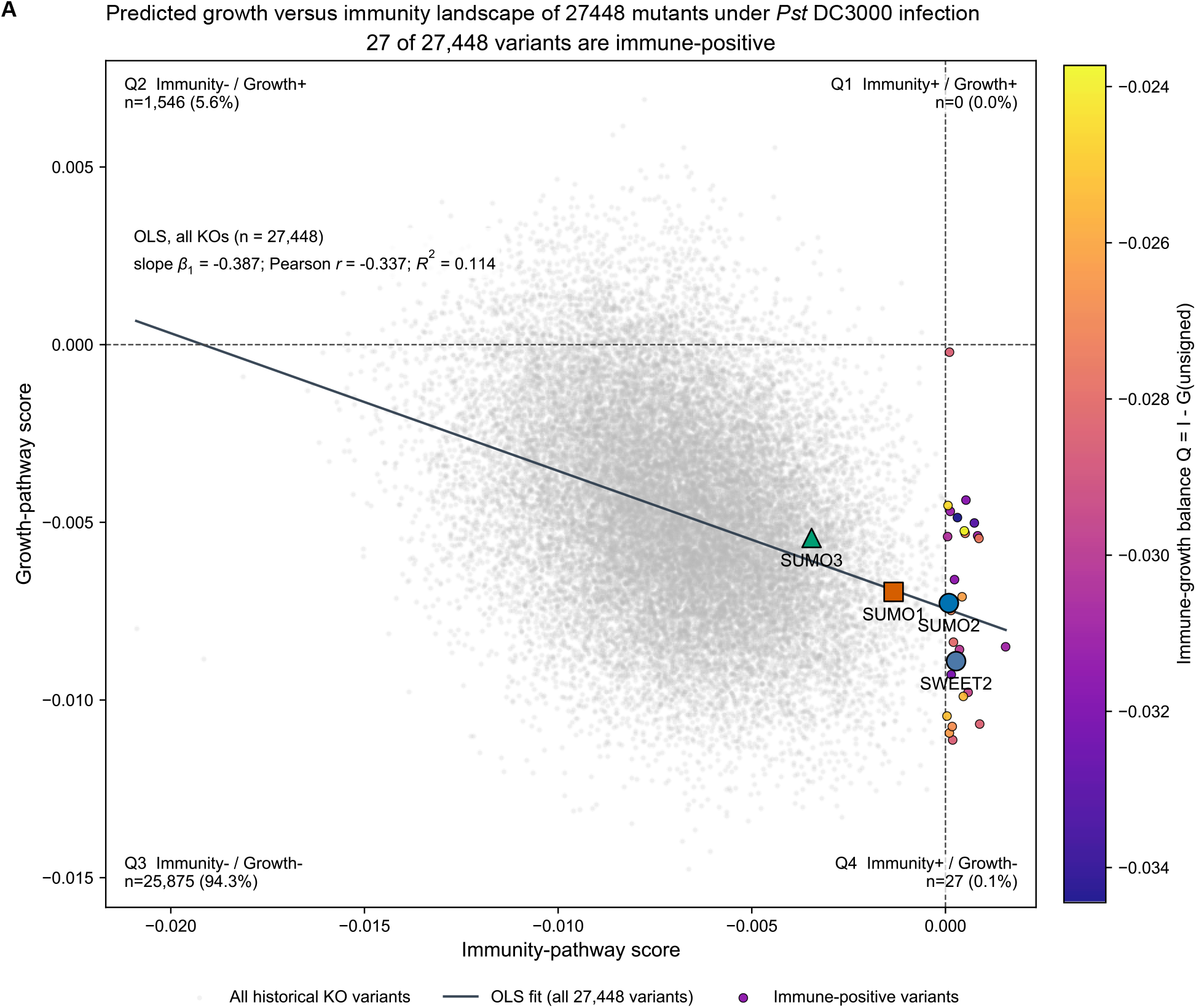
Predicted growth versus immunity landscape of 27,448 mutants under *Pst* DC3000 infection. Each point represents a virtual knockout. The x- and y-axes show the signed immunity-pathway and growth-pathway scores relative to WT, respectively, with dashed lines indicating zero. Gray points represent all 27,448 mutants. The 27 mutants with positive immunity-pathway scores are colored according to the immunity-growth balance score, *Q* = *I − G*, where *G* is the unsigned growth-disruption score. The solid line represents an ordinary least-squares (OLS) regression fitted across all 27,448 mutants (slope *β*_1_ = *−* 0.387, Pearson *r* = *−* 0.337, and *R*^2^ = 0.114), indicating an overall negative association between the predicted immunity- and growth-pathway scores. *SUMO1, SUMO2*, and *SUMO3* are highlighted by an orange square, blue circle, and green triangle, respectively, and *SWEET2* is labeled for reference. *SUMO2* lies on the immunity-positive side, whereas *SUMO1* and *SUMO3* lie on the immunity-negative side. All three have negative signed growth-pathway scores.

**Table S1:** Across complementary sample-wise and gene-wise performance evaluations, the DigiAra hybrid model showed the best overall numerical performance compared with six individual deep- and shallow-learning models and the PerturbMean baseline. Sample-wise performance was assessed using Pearson and Spearman correlations, AUROC, AUPRC, recall of up- and down-regulated genes, mean squared error, and top-1,000 gene overlap. Gene-wise performance was additionally assessed using Pearson correlations calculated across the 34 test samples for each of 27,448 genes.

**Panel A. Correlation and classification performance assessed by Pearson and Spearman correlations, AUROC, and AUPRC.**
| Model | Pearson | Spearman | AUROC | AUPRC |
| --- | --- | --- | --- | --- |
| DigiAra (Hybrid model) | **0.4869 ± 0.2031** | **0.3868 ± 0.1622** | 0.7348 ± 0.0403 | 0.4784 ± 0.1542 |
| Bagging | 0.4479 ± 0.2441 | 0.3437 ± 0.2008 | **0.7389 ± 0.0403** | **0.4788 ± 0.1541** |
| Random forest | 0.4662 ± 0.2277 | 0.3687 ± 0.1760 | 0.7332 ± 0.0351 | 0.4715 ± 0.1470 |
| ExtraTrees | 0.4510 ± 0.2422 | 0.3517 ± 0.1876 | 0.7346 ± 0.0407 | 0.4728 ± 0.1477 |
| XGBoost | 0.4155 ± 0.2200 | 0.3499 ± 0.1680 | 0.7214 ± 0.0384 | 0.4542 ± 0.1512 |
| LightGBM | 0.4510 ± 0.1934 | 0.3728 ± 0.1473 | 0.7279 ± 0.0352 | 0.4665 ± 0.1548 |
| Transformer | 0.3548 ± 0.2294 | 0.2681 ± 0.1561 | 0.6948 ± 0.0685 | 0.4699 ± 0.1530 |
| PerturbMean | 0.3026 ± 0.2809 | 0.2421 ± 0.2249 | 0.7086 ± 0.0450 | 0.4384 ± 0.1643 |

| Model | Rec_Up | Rec_Down | MSE | Overlap (Top 1000) |
| --- | --- | --- | --- | --- |
| DigiAra (Hybrid model) | **0.4025 ± 0.1473** | **0.3901 ± 0.1405** | **0.0808 ± 0.0655** | **0.4110 ± 0.1794** |
| Bagging | 0.3924 ± 0.1438 | 0.3684 ± 0.1481 | 0.0856 ± 0.0651 | 0.3946 ± 0.1996 |
| Random forest | 0.3950 ± 0.1407 | 0.3776 ± 0.1325 | 0.0819 ± 0.0660 | 0.4014 ± 0.1959 |
| ExtraTrees | 0.3919 ± 0.1458 | 0.3696 ± 0.1410 | 0.0842 ± 0.0632 | 0.3994 ± 0.2028 |
| XGBoost | 0.3750 ± 0.1547 | 0.3590 ± 0.1336 | 0.0882 ± 0.0711 | 0.3520 ± 0.1804 |
| LightGBM | 0.3886 ± 0.1421 | 0.3717 ± 0.1344 | 0.0855 ± 0.0671 | 0.3670 ± 0.1640 |
| Transformer | 0.3628 ± 0.1502 | 0.3519 ± 0.1163 | 0.0949 ± 0.0845 | 0.3071 ± 0.2091 |
| PerturbMean | 0.3373 ± 0.1714 | 0.3152 ± 0.1493 | 0.1023 ± 0.0785 | 0.3005 ± 0.1913 |

*Note:* For Panel A and B, values are presented as mean ± standard deviation (SD) across 34 test samples. Boldface indicates the numerically best-performing model for each metric. Higher values indicate better performance for all metrics except mean squared error (MSE), for which lower values indicate better performance. AUROC, area under the receiver operating characteristic curve; AUPRC, area under the precision-recall curve.

**Panel C. Gene-wise prediction performance assessed by Pearson correlation.**
| Model | Gene-wise Pearson |
| --- | --- |
| DigiAra (Hybrid model) | **0.317225** |
| Bagging | 0.286695 |
| Random forest | 0.297848 |
| ExtraTrees | 0.278219 |
| XGBoost | 0.264395 |
| LightGBM | 0.296011 |
| Transformer | 0.188415 |

*Note:* For Panel C, Pearson correlations were calculated across the 34 test samples separately for each of 27,448 genes, and the reported value represents the mean correlation across genes. Boldface indicates the numerically best-performing model for each metric.

**Table S2:** Focused KEGG pathways and their roles in the signed EvaKO Model A analysis.

| KEGG ID | Pathway | Group | Submodule | Analysis role | Weight | KEGG genes, *n* | Notes |
| --- | --- | --- | --- | --- | --- | --- | --- |
| ath04626 | Plant–pathogen interaction | Immunity | Immune signaling | Immunity score (*I*) | 1/6 | 217 | Included as a core plant immune-recognition and response pathway. |
| ath04016 | MAPK signaling pathway – plant | Immunity | Immune signaling | Immunity score (*I*) | 1/6 | 146 | Included as a core plant immune-signal-transduction pathway. |
| ath00940 | Phenylpropanoid biosynthesis | Immunity | Defense metabolism | Immunity score (*I*) | 1/6 | 132 | Included to represent defense-associated phenylpropanoid metabolism. |
| ath00966 | Glucosinolate biosynthesis | Immunity | Defense metabolism | Immunity score (*I*) | 1/6 | 26 | Included to represent glucosinolate-based defense metabolism. |
| ath00380 | Tryptophan metabolism | Immunity | Defense metabolism | Immunity score (*I*) | 1/6 | 65 | Included to represent tryptophan-derived defense metabolism. |
| ath00480 | Glutathione metabolism | Immunity | Defense metabolism | Immunity score (*I*) | 1/6 | 102 | Included to represent redox regulation and defense-associated detoxification. |
| ath04075 | Plant hormone signaling | Plant hormone signaling | – | Context only | – | 433 | Reported as broad hormone context, including salicylic-acid and other hormone branches; not included in *I*, *G*, or *Q*. |
| ath00195 | Photosynthesis | Growth | Photosynthesis & carbon assimilation | Growth penalty (*G*) | 1/11 | 77 | Included to represent the core photosynthetic machinery. |
| ath00196 | Photosynthesis – antenna proteins | Growth | Photosynthesis & carbon assimilation | Growth penalty (*G*) | 1/11 | 22 | Included to represent photosynthetic light-harvesting capacity. |
| ath00710 | Carbon fixation in photosynthetic organisms | Growth | Photosynthesis & carbon assimilation | Growth penalty (*G*) | 1/11 | 71 | Included to represent photosynthetic carbon assimilation. |
| ath01200 | Carbon metabolism | Growth | Central carbon & energy | Growth penalty (*G*) | 1/11 | 275 | Included to represent central carbon metabolism. |
| ath00500 | Starch and sucrose metabolism | Growth | Central carbon & energy | Growth penalty (*G*) | 1/11 | 181 | Included to represent carbohydrate production, storage, and allocation. |
| ath00010 | Glycolysis / Gluconeogenesis | Growth | Central carbon & energy | Growth penalty (*G*) | 1/11 | 120 | Included to represent primary carbon and energy metabolism. |
| ath00020 | Citrate cycle (TCA cycle) | Growth | Central carbon & energy | Growth penalty (*G*) | 1/11 | 64 | Included to represent mitochondrial central energy metabolism. |
| ath00190 | Oxidative phosphorylation | Growth | Central carbon & energy | Growth penalty (*G*) | 1/11 | 169 | Included to represent cellular energy production. |
| ath03010 | Ribosome | Growth | Biosynthesis & replication | Growth penalty (*G*) | 1/11 | 394 | Included to represent protein-synthesis capacity. |
| ath03008 | Ribosome biogenesis in eukaryotes | Growth | Biosynthesis & replication | Growth penalty (*G*) | 1/11 | 98 | Included to represent ribosome assembly and translational capacity. |
| ath03030 | DNA replication | Growth | Biosynthesis & replication | Growth penalty (*G*) | 1/11 | 43 | Included to represent genome replication and proliferative capacity. |

*Notes:* Pathways are listed in the same order used for the focused-pathway heatmap. For each virtual knockout, *I* is the equal-weighted mean Signed-RMS across the six Immunity pathways, whereas *G* is the equal-weighted mean unsigned RMS across the eleven Growth pathways. Plant hormone signaling is reported as biological context and is not used for candidate selection, balance-score calculation, or ranking. Submodules determine grouping and display order only and do not introduce an additional averaging step. The focused analysis contains 18 retained pathways: 6 Immunity, 1 hormone-context, and 11 Growth pathways. The “KEGG genes, *n*” column reports the number of pathway–gene links in the supplied KEGG gene-set summary, obtained from the KEGG REST [link](https://rest.kegg.jp/link/ath/ath04626) endpoint for each pathway. The member genes for a given pathway can be retrieved by replacing the pathway identifier in the endpoint; for example, the genes assigned to ath04626 are available at https://rest.kegg.jp/link/ath/ath04626. These values represent KEGG pathway membership before mapping to the expression index and may therefore differ from the number of genes actually used for scoring.

